# DNA double helix, a tiny electromotor

**DOI:** 10.1101/2022.06.13.495958

**Authors:** Christopher Maffeo, Lauren Quednau, James Wilson, Aleksei Aksimentiev

**Affiliations:** Department of Physics; University of Illinois at Urbana–Champaign; Urbana, IL 61801, USA; Beckman Institute for Advanced Science and Technology, University of Illinois at Urbana–Champaign; Urbana, IL 61801, USA; Department of Bioengineering, University of Illinois at Urbana–Champaign; Urbana, IL 61801, USA

## Abstract

Flowing fluid past curved objects has been used for centuries to power rotary motion in man-made machines. In contrast, rotary motion in nanoscale biological^1, 2^ or chemical^3, 4^ systems is produced by biasing Brownian motion through cyclic chemical reactions.^5, 6^ Here, we show that a curved biological molecule, a DNA or RNA duplex, rotates, unidirectionally, billions of revolutions per minute when electric field is applied along the duplex, with the rotation direction being determined by the duplex chirality. The rotation is found to be powered by the drag force of the electro-osmotic flow, realizing the operating principle of a macroscopic turbine at the nanoscale. The resulting torques are sufficient to power rotation of nanoscale beads and rods, offering an engineering principle for constructing nanoscale systems powered by electric field.

---

While incremental miniaturization of a man-made machine is possible by simply reducing the size of its parts, radical miniaturization typically requires re-evaluation of physical principles that govern the machine’s operation.^7^ For example, in a conventional electromotor, electric field is transformed into rotary motion by means of electromagnetic induction. However, already at the sub-millimeter scale, rotary motion is best produced using electrostatic actuation.^8^ At the nanoscale, biological molecular motors operate with high precision and efficiency^5^ using a chemical reaction to bias direction of random displacement.^6^ Some molecular motors, such as FoF1 ATP synthase^1^ and the bacterial flagellum motor,^2^ are true electromotors, transforming the energy of a transmembrane electric potential into rotation.^9^ Although the biased diffusion mechanism has been realized in purely synthetic molecular systems,^3, 4^ none of those matched the precision and efficiency of a biological motor, which ultimately can be attributed to the latter having structures optimized by evolution.

The years of evolution that furnished DNA might have given us more than just the carrier of the genetic code. The programmable self-assembly of DNA molecules has emerged as a powerful tool for soft nanotechnology,^10, 11^ enabling fabrication of a diverse range of systems.^12–15^ Using the DNA hybridization reaction as fuel,^16^ molecular motors have been constructed to walk,^17–21^ roll,^22^ or pivot^23^ on a pre-defined track, to transport^24^ and sort^25^ molecular cargos, and to undergo reversible conformational transitions.^26^ Aimed at reproducing biological function of molecular motors, multi-subunit concentric DNA origami structures were synthesized to undergo rotary diffusion, driven by stochastic forces.^27–29^ Unidirectional rotation of a self-assembled DNA arm was realized by coupling the arm’s orientation to the direction of the fluid flow and alternating the flow direction in a cyclic pattern.^30^ Yet, it has not escaped our notice that the very shape of a DNA molecule—the shape of a screw—could allow it to function as the simplest possible electromotor.

To investigate if a single DNA duplex will rotate unidirectionally in an external electric field, we constructed an all-atom model of a 16 base pair (bp) DNA duplex surrounded by 1 M KCl electrolyte solution, Fig. 1a. Using the all-atom molecular dynamics method, we simulated the behavior of the duplex when an electric field was applied along its helical axis. The phosphorus atoms of the duplex were restrained to remain at the surface of a cylinder such that the DNA duplex was free to rotate about its axis (see Materials and Methods for technical details).

**Figure 1:**
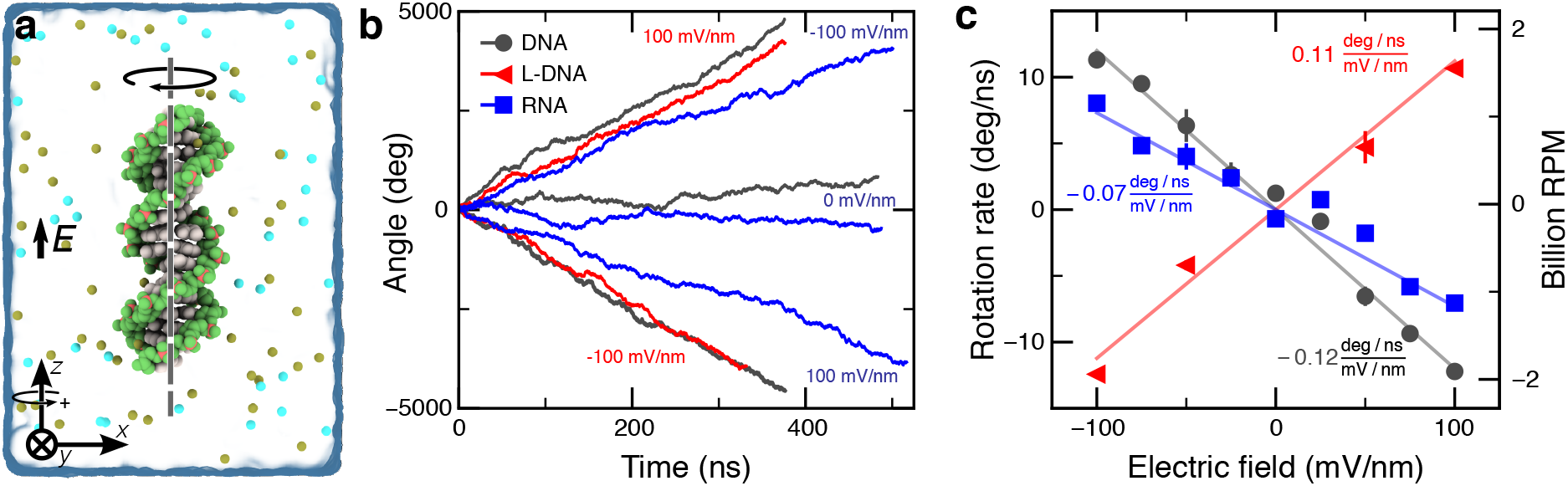
Unidirectional rotation of DNA and RNA molecules in external electric field. **a**, Simulation system containing a 16 bp DNA helix (light gray; green backbone) submerged in 1 M KCl electrolyte (semi-transparent molecular surface); only a fraction of ions is shown. Electric field *E* is applied parallel to the DNA helix. Phosphorus atoms of the DNA are harmonically restrained to the surface of a cylinder such that the DNA is free to rotate about its axis without drifting in the applied field. The circular arrow above the *z* axis indicates a positive rotation direction. The arrow above the DNA indicates the direction of DNA rotation commensurate with the positive direction of the applied field. **b**, Angular displacement of a DNA (black), L-DNA (red) and RNA (blue) helix as a function of simulation time. The sign of displacement is defined in panel a. The electric field strength is annotated near each curve; the purple labels apply to both DNA and RNA. **c**, Average angular velocity of the DNA, L-DNA and RNA helices versus electric field strength. Each data point indicates an average value from a ~400 ns MD trajectory with error bars showing the standard error using 40-ns block-averaged data. Lines depict linear regression fits to the data. The right axis displays the rotational velocity in revolutions per minute (RPM).

With a 100 mV/nm electric field directed along the helical axis, the DNA duplex was observed to rotate about its axis with the rotation vector pointing against the applied field, Fig. 1b and Supplementary Movie 1. Reversing the direction of the applied field reversed the direction of the DNA rotation. In the absence of the field, the DNA duplex was observed to rotate stochastically. In a set of control computational experiments, we repeated our simulations using a 16 bp left-handed duplex constructed from L-DNA, the mirror image stereoisomer of biological DNA, finding the left-handed duplex to rotate with the same velocity as the canonical DNA duplex, but in the opposite direction, Fig. 1b and Supplementary Movie 2. Finally, we repeated the simulation using a 16 bp A-form RNA duplex, Supplementary Movie 3, finding the RNA to rotate in the same direction as the canonical B-DNA duplex, albeit with a reduced rate of rotation.

Repeating the simulations for several values of the electric field yielded the dependence of the average rotation rate on the electric field strength, Fig. 1c. Under a 100 mV/nm electric field—a magnitude well within reach of nanopore translocation experiments^31^—the nucleic acid duplexes were found to spin about 1 billion revolutions per minute, surpassing the rotation rate of the fastest known molecular rotors by several orders of magnitude.

To determine the torque imparted on the nucleic acid structures by the applied electric field and the torque generation mechanism, we built all-atom systems where a DNA or an RNA duplex was connected to itself across the periodic boundary of the simulation system, reproducing the case of an effectively infinite, straight duplex, Fig. 2a. Each phosphorus atom in the duplex was held to its initial coordinates using a harmonic restraining force, Fig. 2b and Supplementary Fig. 1a. Subject to external electric field, equilibrium displacement of the phosphorus atoms relative to their initial coordinates measured the effective forces and torques experienced by the nucleic acid duplex.

**Figure 2:**
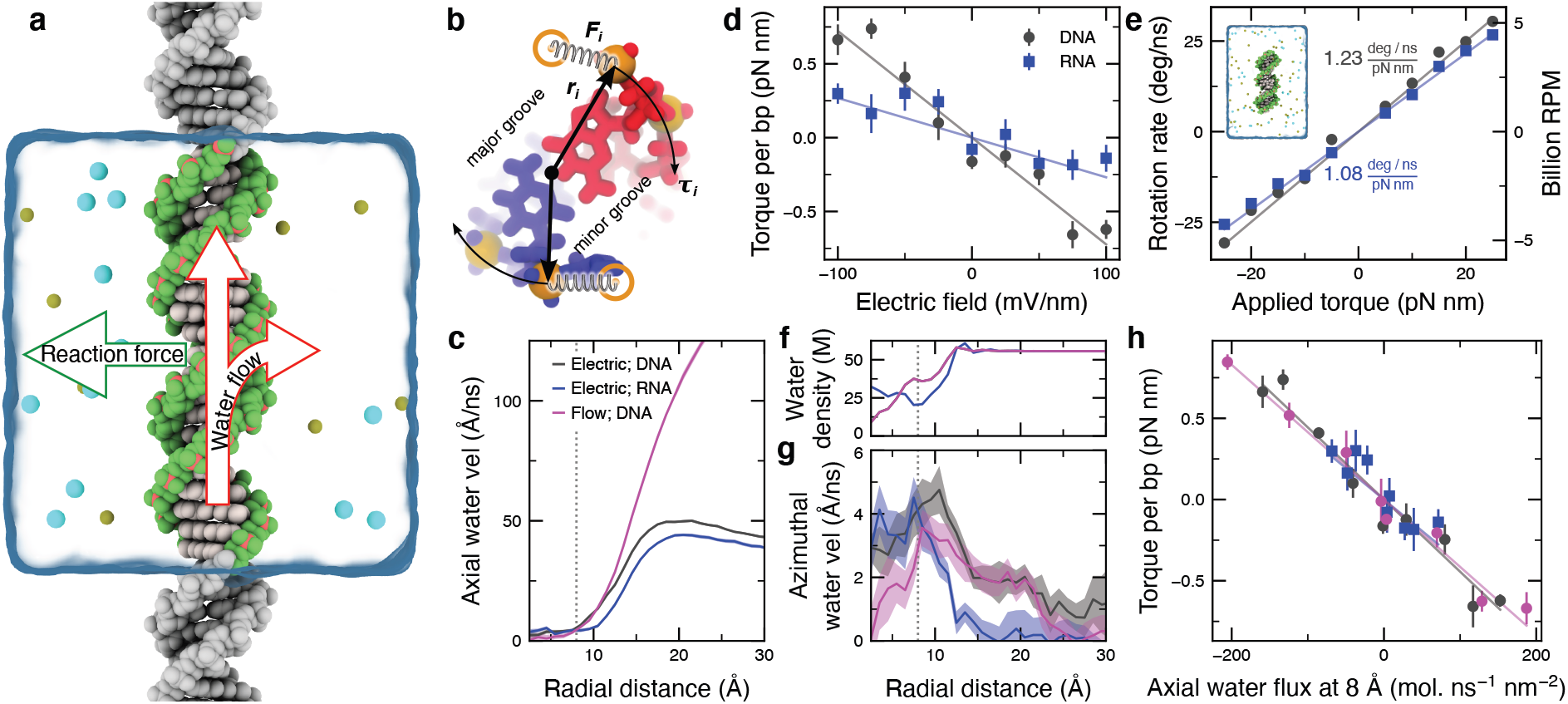
Torque generation mechanism. **a**, Schematics of the simulation system where the DNA was made effectively infinite by connecting its stands over the periodic boundary. The arrows illustrate the torque generation mechanism: redirection of the water flow at the DNA surface. **b**, Illustration of a method used to measure or apply torque. Cross-section of the DNA construct is shown. **c**, Average water velocity along duplex axis versus distance from the axis. Data were averaged over multiple trajectories at ±50 mV/nm electric field or ±12.7 bar/nm pressure gradient using 5 Å radial bins. **d**, Torque per base pair exerted on DNA (black) and RNA (blue) duplex as a function of electric field. Lines show linear fit to the data. Each data point indicates the average value from a ~ 100 ns MD trajectory; the error bars show the standard error computed using 20-ns block-averaged data. **e**, Rotation rate as a function of applied torque. Each data point indicates the average value from a 100–300 ns MD trajectory; the error bars show the standard error computed using 20-ns block-averaged data. The data were obtained using finite length (16 bp) DNA and RNA molecules, one such system is shown in the background. The slope of the linear fit (lines) to the data is the rotational mobility of the duplexes. **f**, Density of water molecules versus distance from the duplex axis. For the DNA system, the electric field and solvent flow data overlap almost exactly. **g**, Average tangential water velocity versus radial distance from the center of the DNA or RNA duplex. Data was computed as described in panel e. The shaded region depicts the magnitude of the difference between data obtained at opposite orientations of the electric field; the solid lines depict the average over the two orientations. **h**, Torque versus axial water velocity 8 Å away from the center of DNA and RNA duplexes.

Similar to our previous simulations of the effective force on a DNA duplex,^32^ we find the effective force to be considerably reduced compared to the force expected from the nominal charge of the DNA molecule, Supplementary Fig. 1b. Previously, we showed that such an effective reduction of the DNA charge is caused, in part, by the drag of the electro-osmotic flow that in turn is produced by the motion of counterions near the DNA surface.^32^ Interestingly, we found the effective force on the RNA duplex to be about 80% of the effective force experienced by a DNA duplex under identical simulation conditions, which we attribute to differences in the fluid flow profiles near the surface of each molecule, Fig. 2c.

We determined the effective torque by multiplying the harmonic force restraining each phosphorus atom by the distance from the duplex axis to that atom, Fig. 2b. Averaging the torque values over all phosphorus atoms and the simulation trajectories yielded the dependence of the effective torque on the applied electric field, Fig. 2d. Under the same electric field, a DNA duplex was found to experience a greater torque per basepair than an RNA duplex, which we attribute to the different shape of the molecules, with A-form RNA having a larger radius, smaller pitch (length of one turn), and more solvent-accessible center, compared to B-form DNA. The torque, *τ*, and the angular velocity, *ω*, are related by *ω* = *μτ*, where *μ* is the rotational mobility (inverse of the rotational friction coefficient), which we determined by simulating forced rotation of the duplexes in the absence of applied electric field, Fig. 2e. The DNA duplex is found to have a slightly higher rotational mobility than an RNA duplex of the same nucleotide composition, in agreement with the mobility values extracted from the analysis of the rotational diffusion of the duplexes, Supplementary Fig. 2. Thus, under the same electric field, an RNA duplex is expected to rotate slower than a DNA duplex because an RNA duplex generates a lower effective torque and has a lower rotational mobility. Indeed, we find the rotation rate observed in our electric field simulations to be prescribed by the product of independently determined effective torque and rotational mobility, Supplementary Fig. 3.

To determine the physical origin of the torque, we determined the magnitude of the fluid velocity directed along, Fig. 2c, and tangential to, Fig. 2g, the axis of the duplex as well as the local density of the fluid, Fig. 2f. Because of the grooves, the water density does not decrease all the way to zero at radial distances less than those of the DNA backbone (~12 Å). Near and within the DNA duplex, the tangential component of the water flux has small yet statistically significant values, indicating that some of the water flux is redirected tangentially around the DNA in the directions expected from the helical geometry of the DNA duplex, Supplementary Movie 4 and 5. The momentum transfer caused by the redirection of the water flow imparts an effective tangential force, producing a non-zero net torque on the duplex, Fig. 2a.

If the rotation of the duplexes in an electric field is, ultimately, caused by the flow of the solvent, then it should be possible to observe the rotation in a system with a flow, in the absence of the applied electric field. To test this hypothesis, we applied a small force along the DNA axis to every water molecule, Supplementary Fig. 4a, producing a steady-state water flow through the periodic DNA system, Fig. 2c. The flow, directed along the duplex axis, was observed to rotate DNA in the expected left-handed direction, Supplementary Fig. 4b and Supplementary Movie 6. The rotation rate was seen to depend linearly on the hydrostatic pressure gradient, Supplementary Fig. 4c. Despite the net flux of solvent being much larger in the pressure-driven simulation than when induced by the electro-osmotic effect, Fig. 2c, the rotation rates in both cases are of similar magnitudes (cf. Fig. 1c and Supplementary Fig. 4c), which we attribute to the similar variation of the flow profile near the surface of the DNA duplex. In fact, we find that the measured torques to fall on the same master curve when plotted versus the magnitude of axial water flux measured 8 Å away from the center of the nucleic acid duplex, Fig. 2h.

To determine how well the torque generated by the duplex can drive rotation of much larger loads, we consider a situation where a DNA molecule is threaded through a nanopore in a thin membrane. The two strands at one end of the DNA molecule are connected to a spherical load, Fig. 3a, which we consider to be located either away from or in close proximity to the membrane. According to our calculations, a 10 mV/nm field applied over a 5 nm (15 bp) length of a DNA duplex generates a 1 pN nm torque. Using the theoretical expressions for the hydrodynamic drag of a spherical particle (see Methods) and assuming a 1 pN nm torque on the duplex, we find the average rotation rate of the spherical particle to decrease from millions to thousands of RPMs as the particle radius increases from 10 to 40 nm, Fig. 3b. Similar calculations for a rod-like load, Fig. 3c, yields rotation rates that decrease with the length of the rod and are in the range of tens of thousands of RPM for a rod 200 nm in length, Fig. 3d. Since the torque on the DNA is cause by the solvent flow, similar magnitude rotations can be expected when the flow is generated using a salinity gradient.^33^

**Figure 3:**
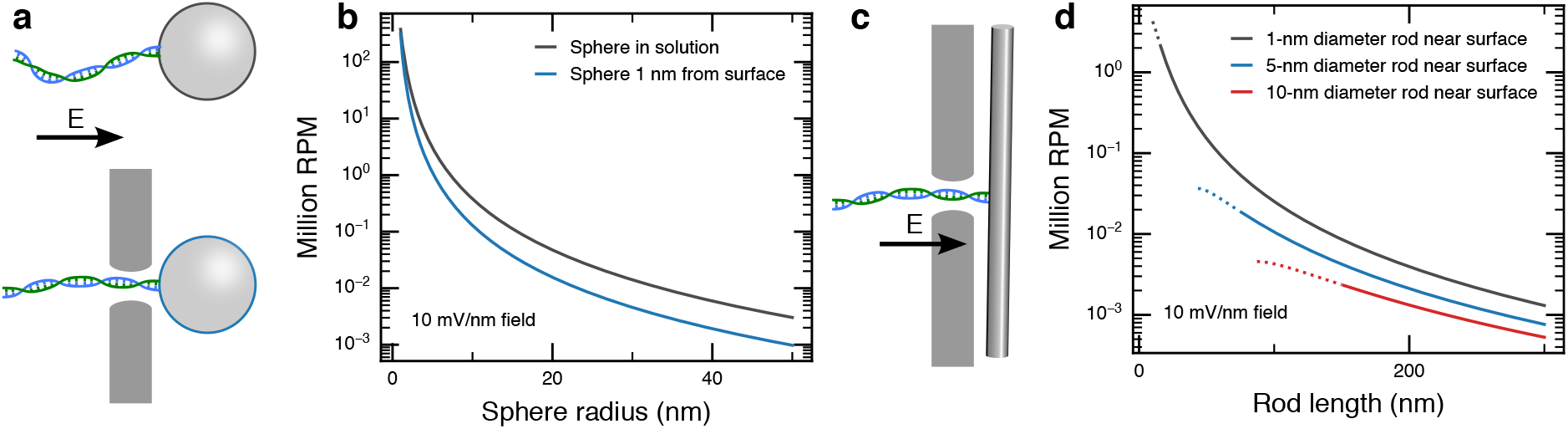
DNA-powered rotation of hydrodynamic loads. **a**, Schematic of a model system where a DNA molecule is tethered, with both of its ends, to a spherical nanoparticle and is threaded through a nanopore in a membrane with the latter being either far from (top, membrane not shown) or near to (bottom) the nanoparticle. Electric field across the membrane generates torque on the DNA. **b**, Theoretical dependence of the rotation rate of a spherical nanoparticle on the nanoparticle’s radius under a 1 pN nm torque on the DNA. **c**, A variant of a model system shown in panel a but with the DNA molecule threaded through a nanopore and tethered to a rod-like nanoparticle. **d**, Theoretical dependence of the rotation rate on the rod length for the system depicted in panel c. Dashed line indicates the regime where the length is comparable to the rod diameter invalidating the approximations used to derive the expression for the rotational diffusion coefficient of a cylinder near a surface.

Our calculations suggest that a DNA molecule is subject to considerable torques in typical nanopore translocation experiments. Given that torsion propagates much faster along a DNA molecule than tension, with the latter being defined by the 3D configuration of the molecule,^34^ the torque applied to the DNA within the nanopore could, potentially, induce the formation of plectonemes in the DNA structure upstream from the nanopore. The passage of plectonemes through a nanopore would register as a transient three-fold increase in the depth of the blockade current, a phenomenon that has been previously attributed to the passage of DNA knots.^35^

In summary, we have shown that the very chiral shape of a DNA duplex is sufficient to generate a rotary motion when solvent moves past its surface. Such fluid motion can be conveniently produced by applying electric field along the DNA helix, utilizing the electro-kinetic effect. The torques generated by the effect are sufficiently large to power rotation of 10–200 nm scale loads with rates on par or exceeding those of biological molecular motors. In contrast to molecular motors, however, the torque generation mechanism in the DNA turbine closely resembles that for a macroscopic turbine, suggesting that other macroscopic machines that rely on fluid flow for their operation can be scaled down to the nanoscale.

## Acknowledgments

CM and AA acknowledge illuminating discussions with Xin Shi, Cees Dekker and Hendrik Dietz.

## Authors contributions

Conceptualization: AA

Methodology: AA, CM, JW

Investigation: CM, LQ, JW

Visualization: CM, LQ, JW

Funding acquisition: AA, CM, LQ, JW

Project administration: AA

Supervision: AA

Writing – original draft: AA, CM, LQ, JW

Writing – review & editing: AA, CM

## Data and materials availability

All data, simulation trajectories and analysis code are available upon request.

