## Supplementary Information for "DNA double helix, a tiny electromotor"

### Supplementary Material
### DNA double helix, a tiny electromotor

Christopher Maffeo,[1,2], Lauren Quednau[3], James Wilson[1], Aleksei Aksimentiev[1,2,3*]

[1]Department of Physics; University of Illinois at Urbana–Champaign; Urbana, IL 61801, USA.

[2]Beckman Institute for Advanced Science and Technology, University of Illinois at Urbana–Champaign; Urbana, IL 61801, USA.

[3]Department of Bioengineering, University of Illinois at Urbana–Champaign; Urbana, IL 61801, USA.

**This PDF file includes:**

Materials and Methods

Supplementary Figure 1 to 4

Captions for Supplementary Movie 1 to 6

**Other Supplementary Materials for this manuscript include the following:**

Supplementary Movie 1 to 6

### Materials and Methods

All MD simulations were performed using the NAMD program,[1] CHARMM36 parameters for DNA, water and ions[2,3] with CUFIX[4] corrections, periodic boundary conditions, and the TIP3P model of water.[5] The long-range electrostatic interactions were computed using the particle-mesh Ewald scheme over a 1 Å-spaced grid.[6] Van der Waals and short-range electrostatic forces were evaluated using the 10–12 Å smooth cutoff scheme. The integration time step was 2 fs, and the full electrostatics were calculated every three time steps.

Five all-atom models of nucleic acid systems were constructed using the nucleic acid builder.[7] The first two models were built from either DNA or RNA nucleotides according to the $(AT)_8$ sequence. A custom conversion script was used to transform the coordinates of the 16-bp DNA fragment to produce a complementary L-DNA model. The two-turn periodic DNA and RNA constructs were built from the sequences $d(ACTG)_5dA$ (10.5 bp per turn) and $(ACUG)_5AC$ (11 bp per turn), respectively. The strands of the periodic constructs were connected to themselves over the periodic boundaries using the LKNA patch.

Water and ions were added to each system using the Solvate and Autoionize plugins of VMD,[8] producing a volume of 1 M KCl solution surrounding a nucleic acid. Each system was minimized for 1000 steps, while harmonically restraining each phosphorus atom to its initial coordinates using a 0.1 kcal mol$^{-1}$Å$^{-2}$ spring constant. The systems were equilibrated for 15 ns under the same position restrains using the Nosé-Hoover Langevin piston with 295 K and 1 atm targets for temperature and pressure, respectively. For non-periodic systems, during the equilibration, the dimensions orthogonal to the DNA axis were kept fixed while the dimension parallel to DNA axis ($z$ axis) was allowed to fluctuate to meet the pressure target. For periodic systems, the orthogonal dimensions could fluctuate independently from the parallel dimension in such a way that the aspect ratio of the orthogonal plane was preserved. The langevin damping coefficient was 0.1 ps$^{-1}$.

All applied electric field simulations were carried out in a constant number of atoms, volume, and temperature ensemble. The system's dimensions were set to the average values from the equilibration. A custom TclForces script was used to harmonically hold each phosphorus atom in DNA or RNA to a cylindrical surface (9.4 Å radius; 0.1 kcal mol$^{-1}$Å$^{-2}$ spring constant).[9] The restraints allow the DNA and RNA molecules to rotate freely about their axes, but restrict them from moving off of a central axis. The phosphorus atoms were also restrained (with the same spring constants) to their idealized positions along the $z$ axis to prevent axial translation of the duplex in external electric field. To prevent fraying, the non-hydrogen atoms forming the terminal base pairs were reinforced (2.9 Å rest length; 4 kcal $\mathrm{mol}^{-1}$Å$^{-2}$ spring constant) using the extrabonds feature of NAMD. The simulations of nucleic acid constructs under a fluid flow were performed using the same protocols as in our applied electric field simulations, except that, instead of an electric field, a constant force directed along the $z$ axis was applied to each water oxygen atom.[10] The resulting flow velocity was determined by the force magnitude, which had a value between 20 to 300 fN for the simulations described in this work.

To measure the force and torque, the simulations were performed exactly as described above, except that the periodic systems were utilized, no cylinder restraints were used, the phosphorus atoms of the duplex were harmonically restrained to their idealized positions (0.1 kcal $\mathrm{mol}^{-1}$Å$^{-2}$ spring constant) to prevent rotation, and frames were written every 120 steps. The torque was calculated at each frame as the sum of the individual torques on the phosphorus atoms $\tau = \sum_i (\vec{F_i} \times \vec{r_i}) \cdot \hat{z}$, where $\vec{r_i}$ is the vector from the center of geometry of the phosphorus atoms to the $i$th phosphorus atom, $\vec{F_i}$ is the restraining force, and $\hat{z}$ is the unit vector along the DNA axis. The forces and torques were determined by post-processing the simulation trajectories with a custom script and were averaged over all frames. Finally, for simulations where a torque was applied to the duplex to cause a rotation, the cylinder restraint TclForces script was modified to additionally apply a constant torque to each phosphorus atom.

The rotation rate calculation for a spherical load particle located either away from or in close proximity to a membrane were performed using the following expressions for the hydrodynamic drag (friction coefficient) of the particle: $\xi_{\mathrm{sph,away}} = 8\pi\eta R^3$[11] and $\xi_{\mathrm{sph,near}} \approx 8\pi\eta R^3(\zeta(3) - 3(\frac{\pi^2}{6} - 1)(\frac{d}{R} - 1))$,[11] respectively, where $\eta = 1.0016$ mPa s is the viscosity of water, $R$ is the radius of the spherical particle, $\zeta(3) \approx 1.20206$ is Riemann's $\zeta$ function evaluated at 3, and $d$ is the distance from the center of the sphere to the membrane. The rotation rate calculations for a rod-like load were performed using the following expression for the friction coefficient of the rod: $\xi_{\mathrm{rod}} = \frac{8}{3}\pi\eta L^3/(\ln(\frac{L}{2R}) - 0.447)$,[12] where $L$ is half the length and $R$ is the radius of the rod. In all cases, the rotation rate was computed as $\omega_{\mathrm{particle}} = \tau/\xi_{\mathrm{particle}}$, where $\tau$ was the effective torque on the duplex and $\xi_{\mathrm{particle}}$ was the friction coefficient of the particle.

#### Supplementary Figures

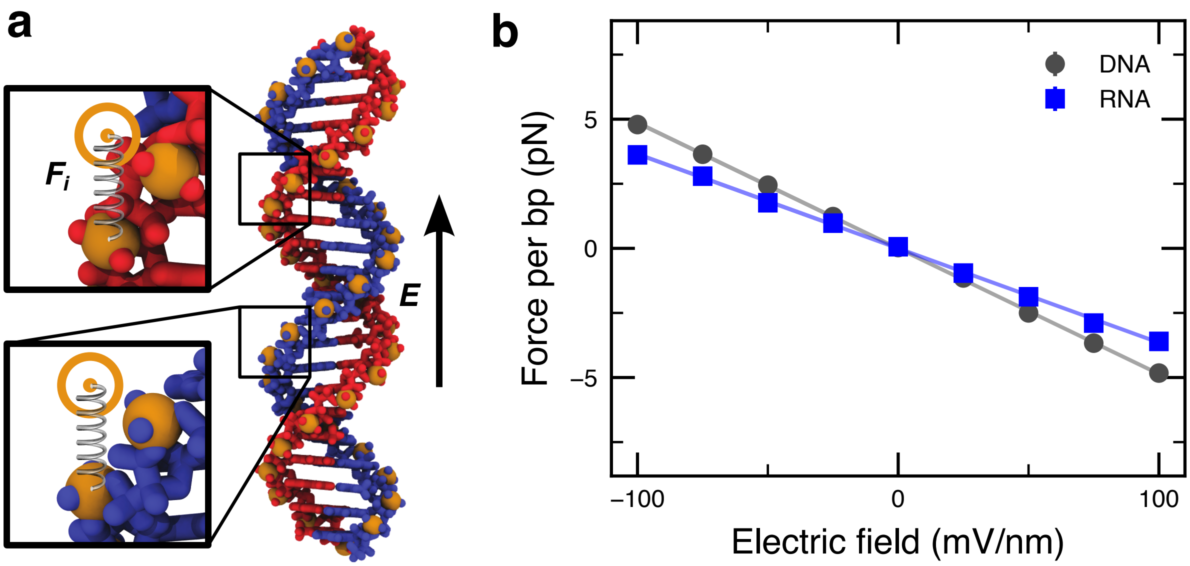


**Supplementary Figure 1: Effective force of the electric field on DNA and RNA molecules**. **a**, Simulation system containing a 21 bp DNA helix (blue and red strands; orange phosphorus atoms) submerged in a volume of 1 M KCl electrolyte (not shown). The insets schematically illustrate how the phosphorus atoms of the molecule are harmonically restrained to their initial coordinates. Equilibrium displacement of the atoms from their initial coordinates multiplied by the spring constant of the harmonic restraint equals by magnitude the effective force experienced by the molecule. **b**, Effective force per base pair of a DNA (black) or an RNA (blue) molecule as a function of the applied electric field.

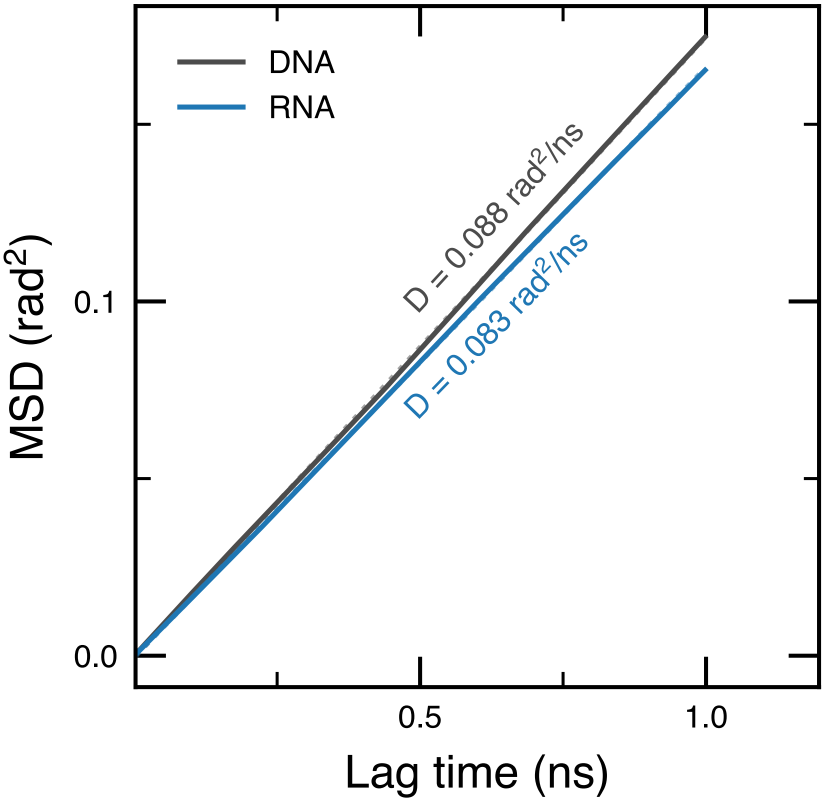


**Supplementary Figure 2: Rotational diffusion of DNA and RNA molecules**. Mean squared angular displacement (MSAD) of a 16-bp DNA (black) and a 16-bp RNA (blue) duplex is plotted versus the lag time. The MSAD values were determined from the analysis of the 500 ns angular displacement traces obtained under zero applied electric field conditions; the traces are shown in Fig. 1 of the main text. Dividing the slopes of the MSAD curves by a factor of two yields the rotational diffusion constants, $D$, of DNA and RNA duplexes at 0.088 and 0.083 $rad^2$/ns, respectively. The corresponding rotational mobilities, $\mu = D/k_BT$, where $k_BT$ is the thermal energy, are $\mu_{\mathrm{DNA}} = 1.24$ deg ns$^{-1}$ pN$^{-1}$ nm$^{-1}$ and $\mu_{\mathrm{RNA}} = 1.17$ deg ns$^{-1}$ pN$^{-1}$ nm$^{-1}$.

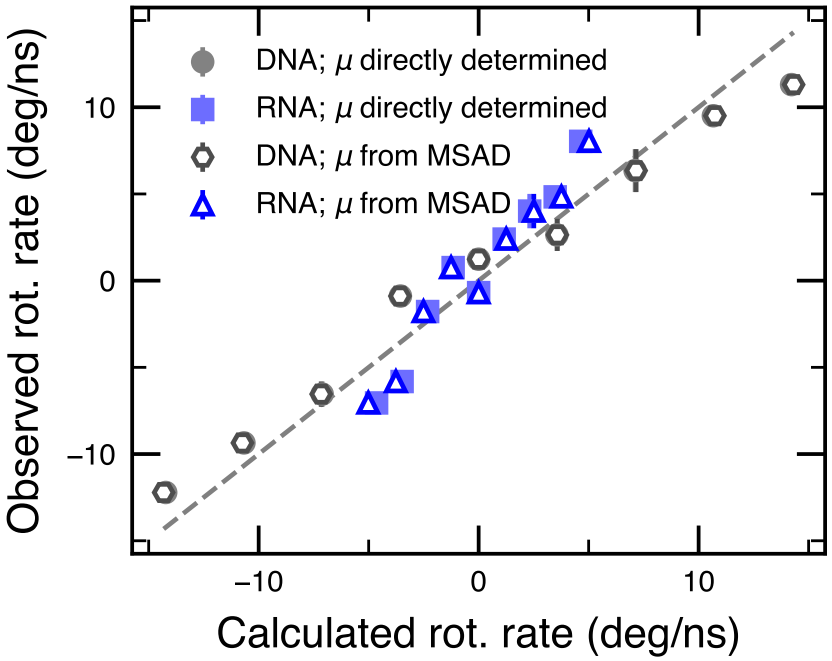


**Supplementary Figure 3: Directly observed versus computed rate of rotation of a finite 16-bp duplex**. The rate of rotation observed in the applied electric field simulations, Fig. 1c of the main text, is plotted versus the rotation rate calculated using two estimates of the angular mobility $\mu$ and the direct measurement of the torque under electric field, Fig. 2d. The torque per base pair was estimated from the applied field strength using the linear regression fit to the data, Fig. 2d. The rotation rate $\omega$ was calculated from the definition of mobility, $\mu = \omega/\tau$, where $\tau$ is the torque on the duplex. Two strategies were used to determine the mobility: using the slope of the mean squared angular displacement (MSAD) as a function of lag time, Supplementary Fig. 2, and directly using the mobility determined from a linear regression fit to rotation rate due to a constant applied torque, Fig. 2e of the main text.

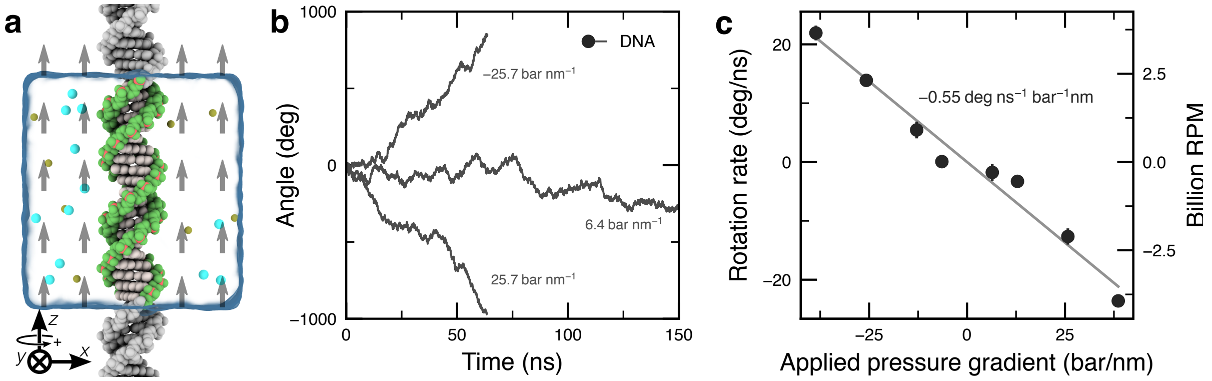


**Supplementary Figure 4: Water flow-induced rotation of a DNA molecule**. **a**, Simulation system containing a 21 bp DNA helix (light gray; green backbone) submerged in a volume of 1 M KCl electrolyte (semi-transparent molecular surface); only a fraction of ions is shown explicitly, for clarity. The DNA molecule is made effectively infinite by connecting each of its strands to itself over the periodic boundary of the simulation unit cell. For illustration, partial periodic images of the DNA molecule are shown in gray. The water flow is produced by applying a small, constant force to each water molecule parallel to the axis of the DNA molecule, generating a pressure differential of $P = Nf/A$, where the force $f$ applied to each of $N$ water molecules in the system, and $A$ is the area of the system within the plane orthogonal to the direction of the applied force.[10] The pressure gradient was then computed by dividing the pressure by the length of the system. Phosphorus atoms of the DNA are harmonically restrained to the surface of a cylinder such that the DNA is free to rotate about its axis; additional restrains prevent the molecules from drifting in the direction of the flow. **b**, Angular displacement of the DNA molecule as a function of simulation time. The sign of displacement is prescribed by the right-hand rule with respect to the positive direction of the force applied to the water molecules, as indicated in panel a. **c**, Average angular velocity of the DNA helix versus pressure gradient due to force applied to each water molecule. Each data point was determined by averaging a ∼60–200 ns MD trajectory. The line shows a linear regression fit to the data. The right axis displays the rotational velocity of the DNA in units of revolutions per minute (RPM).

### Captions to Supplementary Movies

**Supplementary Movie 1.** Rotation of a DNA duplex driven by a 100 mV/nm electric field applied along the duplex axis in the upward direction. The sixteen base pair duplex is shown using vdW spheres colored in light gray (bases) and green (backbone). For clarity, the volume occupied by 1 M KCl electrolyte and the ions are not shown. Phosphorus atoms of the duplex are harmonically restrained to the surface of a cylinder such that the DNA is free to rotate about its axis without drifting in the applied field. The movie illustrates a 60 ns excerpt of a 200 ns MD trajectory.

**Supplementary Movie 2.** Rotation of an L-DNA duplex driven by a 100 mV/nm electric field applied along the duplex axis in the upward direction. The sixteen base pair duplex is shown using vdW spheres colored in light gray (bases) and green (backbone). For clarity, the volume occupied by 1 M KCl electrolyte and the ions are not shown. Phosphorus atoms of the duplex are harmonically restrained to the surface of a cylinder such that the DNA is free to rotate about its axis without drifting in the applied field. The movie illustrates a 60 ns excerpt of a 188 ns MD trajectory.

**Supplementary Movie 3.** Rotation of an RNA duplex driven by a 100 mV/nm electric field applied along the duplex axis in the upward direction. The sixteen base pair duplex is shown using vdW spheres colored in light gray (bases) and green (backbone). For clarity, the volume occupied by 1 M KCl electrolyte and the ions are not shown. Phosphorus atoms of the duplex are harmonically restrained to the surface of a cylinder such that the DNA is free to rotate about its axis without drifting in the applied field. The movie illustrates a 60 ns excerpt of a 199 ns MD trajectory.

**Supplementary Movie 4.** Streamlines representing the average flow of fluid past a periodic DNA duplex when a 100 mV/nm electric field is applied along the duplex' axis in the upward direction. Streamlines are colored from white to red or blue in proportion to the speed with which the fluid moves. The color signifies whether the streamline winds in a left- (blue) or right-handed (red) direction around the duplex. Streamlines were generated by binning the water oxygen atoms into a roughly 1-Å resolution three-dimensional grid and measuring the displacement of each water molecule after a 240 fs interval, taking the mean in each bin over the entire 100 ns trajectory. The resulting grids of water velocities and local densities were passed into the streamline algorithm of the yt[13] Python package, and a custom script was used to write commands for visualization using VMD.[8]

**Supplementary Movie 5.** Streamlines representing the average flow of fluid past a periodic RNA duplex when a 100 mV/nm electric field is applied along the duplex' axis in the upward direction. Streamlines are colored from white to red or blue in proportion to the speed with which the fluid moves. The color signifies whether the streamline winds in a left- (blue) or right-handed (red) direction around the duplex. Streamlines were generated by binning the water oxygen atoms into a roughly 1-Å resolution three-dimensional grid and measuring the displacement of each water molecule after a 240 fs interval, taking the mean in each bin over the entire 100 ns trajectory. The resulting grid of water velocities and local densities were passed into the streamline algorithm of the yt[13] Python package, and a custom script was used to write commands for visualization using VMD.[8]

**Supplementary Movie 6.** Rotation of a DNA duplex under a solvent flow produced by a 12.7 bar/nm hydrostatic pressure gradient. The 21 bp duplex is shown using vdW spheres colored in light gray (bases) and green (backbone). The two strands of the duplex are connected to themselves across the periodic boundary of the simulation unit cell (not shown), making the duplex effectively infinite in the direction of the pressure gradient. For clarity, the volume occupied by 1 M KCl electrolyte and the ions are not shown. Phosphorus atoms of the duplex are harmonically restrained to the surface of a cylinder such that the DNA is free to rotate about its axis without drifting in the solvent flow. The movie illustrates a 60 ns excerpt of a 120 ns MD trajectory.
